# Genetic dissection of rapid proteolysis identifies TXNDC15 as a key factor of ERAD and lipid homeostasis

**DOI:** 10.64898/2026.04.01.715723

**Authors:** Yuyang Liu, Michelle Yinfei Yaochai, Shanshan Liu, Hanan Alwaseem, Kivanc Birsoy

**Affiliations:** Laboratory of Metabolic Regulation and Genetics, The Rockefeller University, New York, NY, USA; The Proteomics Resource Center, The Rockefeller University, New York, NY, USA

## Abstract

Biological systems face a constantly changing environment and must swiftly respond to stimuli, yet how cells sense and adapt to environmental and physiological cues is incompletely understood. Short-lived proteins can be rapidly induced upon perturbation, enabling swift activation of adaptive cellular responses. Leveraging genome-wide data on protein-transcript correlation, we systematically searched for rapid proteolysis substrates whose abundance reflects the activity of the underlying proteolytic machinery. Here, focusing on the candidate substrate ABHD2, we employed CRISPR-based functional screens to dissect its degradation and identified *TXNDC15* as an essential factor in MARCHF6-mediated ER-associated protein degradation (ERAD). Unexpectedly, TXNDC15 supports substrate exit and degradation from the ER via a catalysis-independent mechanism. Loss of *TXNDC15* remodels the ER proteome and lipid homeostasis. Together, our work defines a missing component of ERAD and provides a generalizable strategy to decode post-translational regulatory circuits.

## Introduction

Cells need to respond rapidly to sudden changes in their environment. Post-translational control of protein stability serves as a versatile regulatory mechanism for cells to swiftly adapt to such stimuli^1^. Short-lived proteins are commonly involved in these adaptive responses, given that they can be rapidly derepressed and activated under stress via blockade of their degradation, and timely turned off when the stress is withdrawn. This mode of regulation is especially relevant for compartmentalized organelles, whose internal metabolic environment is not immediately accessible to the nuclear transcriptional machinery^2,3^. Additionally, inter-organellar heterogeneity predicates that organelles will experience localized stimuli or stress that requires a localized adaptive response, most effectively achieved via posttranslational-level regulation.

The endoplasmic reticulum is such an organelle with both a distinct internal environment and a highly plastic composition sensitive to various environmental stimuli and metabolic stress. To function as an essential hub for signaling and metabolism, the ER relies on a dynamically regulated organellar proteome, tightly controlled by a dedicated network of protein machinery in the Endoplasmic Reticulum-Associated Protein Degradation (ERAD) process. Central to this process are more than a dozen of E3 ligases that recognize their client proteins via different adaptors and accessory proteins and safeguard ER function under different nutrient and stress signals^4^. The wide array of substrates that undergo proteolysis via the ERAD process contains numerous components of the canonical nutrient and stress-sensing pathways; the functional complexity of this process implicates that many ERAD factors or substrates likely harbor unknown functions in adaptive signaling.

Given that rapid proteolysis, including ERAD, has been coopted by many mechanisms that respond to environmental stimuli, we leveraged a proteome-wide database for transcript/protein abundance correlation^5^ and searched for unknown regulators of the stability of short-lived proteins in response to metabolic stress. Using this strategy, we uncovered a/b-hydrolase domain containing 2 (ABHD2), a type II membrane protein, as a novel substrate for the lipid-sensitive ERAD E3 ligase MARCHF6. Using this protein as a reporter for a genome-wide CRISPR screen, we identified TXNDC15, a poorly studied protein unknown in ER homeostasis and lipid metabolism, as an essential component of MARCHF6-dependent ERAD. Surprisingly, TXNDC15 supports the degradation of MARCHF6 substrate via a catalytic-independent. The absence of TXNDC15 leads to the rewiring of cellular lipidome reminiscent of those seen in MARCHF6-deficient cells, highlighting the critical role of this pathway in sensing membrane lipid composition and the adaptive maintenance of cellular lipid homeostasis.

## Result

### Systematic analysis of proteome/transcriptome bias identifies ABHD2 as a short-lived nutrient-responsive protein

Given that regulated proteolysis has been co-opted by numerous nutrient and stress-sensing pathways, we hypothesized that proteins with an exceptionally short half-life would be enriched in novel players in nutrient-sensing mechanisms. To search for candidate genes involved in such processes, we systematically analyzed a proteome-wide database of mRNA/protein abundance (*OpenCell* database)^5^ and filtered for proteins on the very low end of protein/mRNA copy number ratios (**Figures 1A and S1A, Data S1**). Among the top 1% of candidate genes, we further narrowed down our search to include proteins with functional annotations related to metabolism and not previously known to undergo rapid proteolysis (**Figure S1A**).

**Figure 1.**
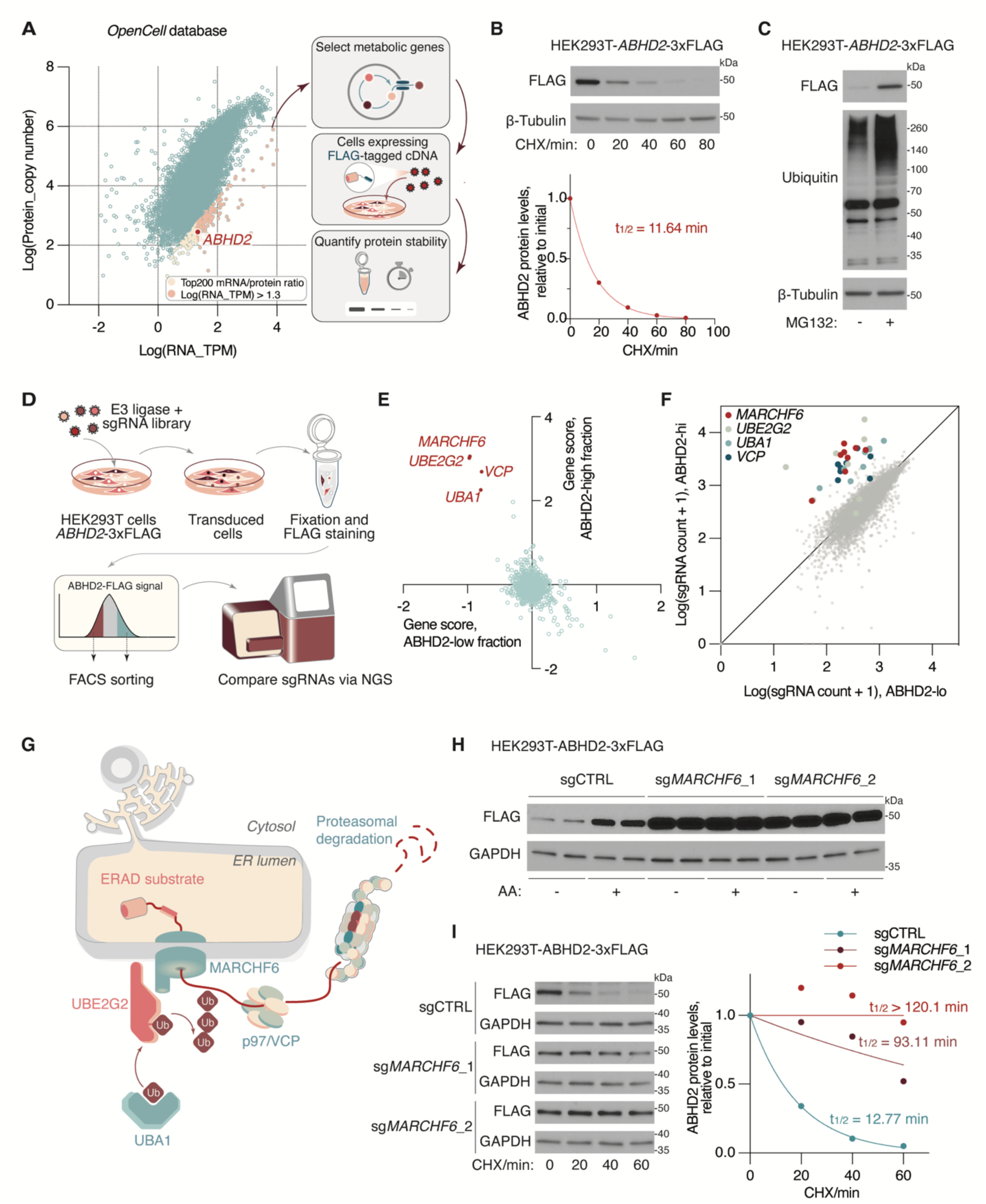
Functional genomic profiling of short-lived metabolic proteins identified ABHD2 as a MARCHF6 target. (**A**) Schematic of the workflow for selecting candidate short-lived metabolic proteins. The dot plot represents proteome-wide measurement of mRNA abundance versus protein copy number, as retrieved from the *OpenCell* database. (**B**) Top, immunoblots of the indicated proteins from HEK293T cells stably expressing *ABHD2*-3xFLAG cDNA. The cells were treated with cycloheximide (CHX, 50 µg/ml) for the indicated time. β-tubulin was used as a loading control. Bottom, quantification of the FLAG bands signal intensity across time. Protein half-life was calculated by the non-linear fitting of FLAG signal intensity versus time to one phase decay exponential model. β-tubulin was used as a loading control. (**C**) Immunoblots of the indicated proteins from HEK293T cells stably expressing *ABHD2*-3xFLAG cDNA, treated with MG132 (25 µM) or DMSO as a control for 4 hours. (**D**) Schematic of the workflow for FACS-based CRISPR screen for factors involved in ABHD2 degradation. (**E**) Scatter plot representing the CRISPR screen result. Points indicate the median relative fold change of sgRNA counts (gene score) of all ubiquitin-proteasome pathway-related genes in the ABHD2-high versus the ABHD2-low fraction. (**F**) Scatter plot showing the distribution of sgRNA abundance in the ABHD-high versus ABHD-low fraction. sgRNAs targeting the selected genes are highlighted in the indicated colors. (**G**) Schematic showing the components of protein machinery involved in MARCHF6-mediated substrate degradation. (**H**) Immunoblots of the indicated proteins from HEK293T cells stably expressing ABHD2-3xFLAG cDNA and the indicated sgRNAs, treated with arachidonic acid (AA, 50 µM) or BSA as a control for 4 hours. GAPDH was used as a loading control. (**I**) Left, degradation kinetics of ABHD2-3xFLAG as shown by immunoblots of the indicated proteins from HEK293T cells stably expressing *ABHD2*-3xFLAG cDNA, treated with cycloheximide (CHX, 50 µg/ml) or DMSO as a control for the indicated time. Cells were stably expressing the indicated sgRNAs. GAPDH was used as a loading control. Right, quantification of the FLAG bands signal intensity across time. Protein half-life was calculated by the non-linear fitting of FLAG signal intensity versus time to one phase decay exponential model.

The candidate ORFs were cloned into a reporter construct, in which the C-terminal 3xFLAG-tagged coding sequence of the candidate protein is co-translated with an HA-tagged RFP as an internal control. We screened the relative stability of candidate proteins by treating HEK293T cells expressing the reporter constructs with the translation inhibitor cycloheximide (CHX) or the proteasome inhibitor MG132 for a short period of time. Among the 22 candidates prioritized for testing, 10 proteins displayed a markedly reduced FLAG/HA ratio upon CHX treatment and an increased FLAG/HA ratio upon MG132 treatment (**Figure S1B**), indicating a proteasome-dependent proteolysis mechanism.

We prioritized our analysis on organellar metabolic enzymes with unknown degradation mechanisms. Therefore, we focused our analysis on alpha/beta hydrolase domain-containing 2 (ABHD2), a lipid metabolism modulator previously implicated in sperm activation yet not known to be subjected to proteolytic regulations^67^. CHX-chase experiment indicated that ABHD2 has a remarkably short half-life (∼ 12 min) and displayed marked stabilization upon inhibition of proteasome function by MG132 (**Figures 1B and 1C**). These results indicate that ABHD2 is subjected to active proteasome-dependent degradation and could potentially be induced upon environmental perturbation, providing a mechanistic basis for a rapidly responsive post-translational response circuit for metabolic adaptation.

### ABHD2 is a target of the MARCHF6 E3 ligase

Given its short half-life, we next sought to harness ABHD2 as a reporter for uncovering novel players in its regulation. To first determine the proteolysis mechanism of ABHD2, we performed a FACS-based CRISPR screen on a stably expressed ABHD2-3xFLAG transgene reporter. We designed a targeted CRISPR sgRNA library covering ∼800 genes annotated to be involved in the ubiquitin-proteasome pathway (**Figure 1D**). Fixed and permeabilized cells were immunostained with fluorophore-conjugated anti-FLAG M2 antibody; the subpopulations of cells displaying the highest or lowest FLAG signals were sorted by FACS, and sgRNA sequences were quantified by next-generation sequencing^8^. Remarkably, the top genes with differential sgRNA enrichment in the ABHD2-hi and -lo populations were the endoplasmic reticulum-resident E3 ligase *MARCHF6*, its cognate E2 ubiquitin-conjugating enzyme *UBE2G2*, the E1 ubiquitin-activating enzyme *UBA1*, and AAA-ATPase p97/*VCP* responsible for the retro-translocation of ER-resident protein (**Figure 1E-1G, Data S2A and S2B**). These results identify ABHD2 as a target for ER-associated protein degradation (ERAD) mediated specifically by MARCHF6.

MARCHF6/Doa10 is a highly conserved multi-pass membrane protein that functions as an E3 ligase for numerous proteins involved in lipid metabolism^9^, whose activity is tuned by membrane lipid composition^10^. Consistent with the lipid-sensitive activity of MARCHF6, ABHD2 protein levels are significantly induced upon treatment with unsaturated, but not saturated fatty acids (**Figure S2A**). Interestingly, ABHD2 is also stabilized by its reported substrate 2-arachidonoylglycerol. A follow-up CRISPR screen with a metabolism-focused sgRNA library yielded top-scoring genes known to be involved in the regulation of membrane lipid saturation (**Figures S2B-S2D, Data S2C**). Confirming the screen results, knocking out *MARCHF6* significantly increased steady-state ABHD2 protein levels, prolonged its half-life and abolished its response to the treatment with polyunsaturated fatty acids (**Figures 1H and 1I**), supporting that ABHD2 is a target of lipid-sensitive ERAD mediated by MARCHF6.

### A genome-wide CRISPR screen identifies TXNDC15 as a novel player in MARCHF6-dependent ERAD

Despite the well-documented role of MARCHF6 in ERAD and lipid homeostasis, the mechanism for its substrate recognition and processing remains incomplete. We therefore hypothesized that ER-localized ABHD2 (**Figure 2A**) could be used as a reporter to identify additional players in this homeostatic mechanism. Therefore, we repeated our CRISPR screen to cover a whole-genome library using the same workflow (**Figure 2B**). Reassuringly, *MARCHF6*, *UBE2G2*, *UBA1* and *VCP* remained near the top of the genes displaying significant enrichment in ABHD2-hi fraction of cells (**Figure 2C and Data S2D**); among the hits of the screen were also subunits of the proteasome and auxiliary/adaptor proteins for p97/VCP (**Figures 2D and S2E, Data S2E**). Unexpectedly, the strongest hit was *TXNDC15*, a poorly characterized gene previously known to be involved in cilia biogenesis (**Figure 2, C to E, Data S2E**). Remarkably, an analysis of gene effects in ∼500 cancer cell lines from Depmap showed high levels of co-essentiality between *MARCHF6* and *TXNDC15* (ranked 1^st^ and 3^rd^, respectively), suggesting a tight functional connection between the two proteins (**Figure 2F**). Reminiscent of MARCHF6, knocking out TXNDC15 also dramatically stabilized ABHD2 and prolonged its half-life (**Figures 2G, 2H and S2F**). These results indicate that TXNDC15 is a key factor mediating MARCHF6-dependent ERAD process.

**Figure 2.**
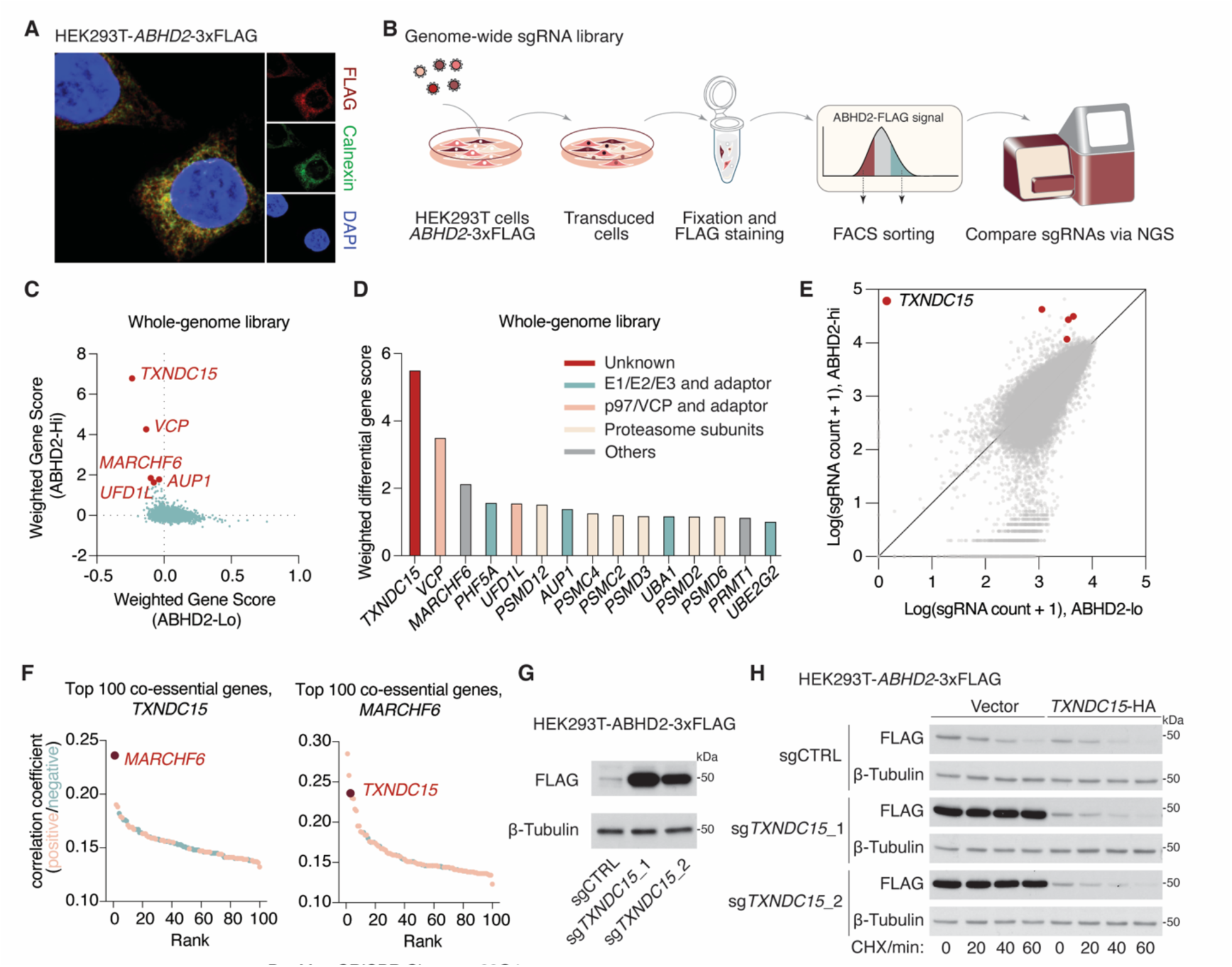
TXNDC15 is a key mediator of MARCHF6-mediated ERAD. (**A**) Immunofluorescence images of ABHD2-3xFLAG (red), Calnexin (green) and DAPI (blue) in HEK293T cells. (**B**) Schematic of the workflow for FACS-based genome-wide CRISPR screen for factors involved in ABHD2 degradation. (**C**) Dot plot representing the CRISPR screen result. Points indicate the sgRNA count-weighted-average relative fold change of sgRNA counts (gene score) of all quantified genes in the ABHD2-high versus the ABHD2-low fraction. (**D**) Weighted differential gene score and functional annotation of top hit genes in the CRISPR screen. (**E**) Scatter plot showing the distribution of sgRNA abundance in the ABHD-high versus ABHD-low fraction. sgRNAs targeting *TXNDC15* are highlighted in red. (**F**) Dot plot representing the top 100 genes ranked by the correlation of their gene effects in over 1,000 cancer cell lines with *MARCHF6* or *TXNDC15*. (**G**) Immunoblots of the indicated proteins from HEK293T cells stably expressing *ABHD2*-3xFLAG cDNA and the indicated sgRNAs. β-tubulin was used as a loading control. (**H**) Degradation kinetics of ABHD2-3xFLAG as shown by immunoblots of the indicated proteins from HEK293T cells stably expressing *ABHD2*-3xFLAG cDNA, treated with cycloheximide (CHX, 50 µg/ml) or DMSO as a control for the indicated time. Cells were stably expressing the indicated sgRNAs and *TXNDC15-*HA cDNA or an empty vector as a control. GAPDH was used as a loading control.

### TXNDC15 assists the degradation of MARCHF6 substrate via a catalytic-independent mechanism

TXNDC15 belongs to the TMX protein family, which harbors membrane-anchoring helices and one or more thioredoxin domains^11^. Immunofluorescence imaging showed that TXNDC15 localizes to the endoplasmic reticulum where MARCHF6 resides (**Figure 3A**). Alpha-fold modeling of protein structure indicated that TXNDC15 harbors a single transmembrane helix at the C-terminus, a long, acidic unstructured region near the N-terminus and a thioredoxin domain in between (**Figure 3B**). To functionally dissect the role of each structural element of TXNDC15, we constructed a series of truncated proteins and tested their ability to interact with MARCHF6. Immunoprecipitation experiments showed that TXNDC15 indeed interacts with MARCHF6 protein (**Figure 3C**). Deletion of the thioredoxin domain caused a quite modest decrease in TXNDC15-MARCHF6 interaction. However, removal of the transmembrane helix and splicing the truncated TXNDC15 with a different ER-anchoring helix from OST4 caused a stronger loss of TXNDC15-MARCHF6 interaction, indicating a significant contribution of this domain to interaction and possibly functional coordination with MARCHF6. Next, we tested the ability of the truncation constructs to restore ABHD2 degradation in TXNDC15-knockout cells. Loss of the thioredoxin domain completely abolished TXNDC15 function in ABHD2 degradation, highlighting its functional essentiality (**Figure 3D**). On the other hand, the removal of the N-terminus unstructured region didn’t seem to severely compromise protein function. Since numerous auxiliary factors in the cytosol also assist the degradation of ER-resident proteins, we examined the function of the short C-terminus cytosolic fragments of TXNDC15 by removing the amino acid residues beyond aa350 of TXNDC15. Although this truncated protein displayed poor expression levels, it still retains intact function in promoting ABHD2 proteolysis (**Figure S3A**).

**Figure 3.**
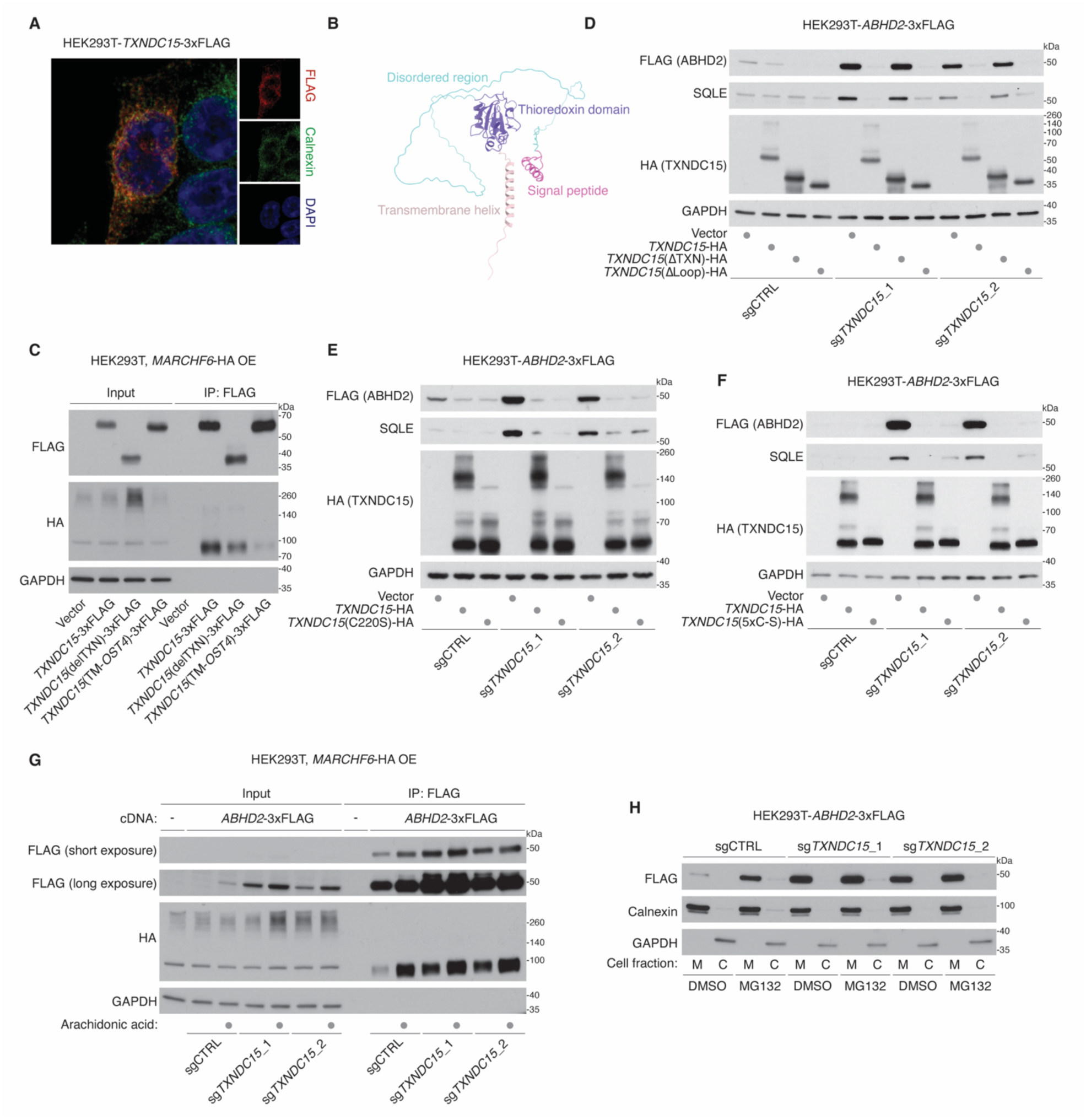
TXNDC15 is a key mediator of MARCHF6-mediated ERAD. (**A**) Immunofluorescence images of TXNDC15-3xFLAG (red), Calnexin (green) and DAPI (blue) in HEK293T cells. (**B**) AlphaFold2-predicted structural model of TXNDC15, with its different structural elements highlighted in the indicated colors. (**C**) Immunoblots showing the indicated proteins in whole-cell lysate or FLAG-immunoprecipitant from HEK293T cells transfected with *MARCHF6-*HA cDNA, along with wild-type or in the indicated modified *TXNDC15-*3xFLAG cDNAs. GAPDH was used as a loading control. (**D**) Immunoblots of the indicated proteins from HEK293T cells stably expressing *ABHD2*-3xFLAG cDNA and the indicated sgRNAs. Cells were complemented with cDNA of wild-type or the indicated truncated constructs of *TXNDC15*-HA. GAPDH was used as a loading control. (**E**) Immunoblots of the indicated proteins from HEK293T cells stably expressing *ABHD2*-3xFLAG cDNA and the indicated sgRNAs. Cells were complemented with cDNA of wild-type or the C220S-mutant *TXNDC15*-HA. GAPDH was used as a loading control. (**F**) Immunoblots of the indicated proteins from HEK293T cells stably expressing *ABHD2*-3xFLAG cDNA and the indicated sgRNAs. Cells were complemented with cDNA of wild-type or the all cysteine-to-serine-mutant (5xC-S) *TXNDC15*-HA. GAPDH was used as a loading control. (**G**) Immunoblots showing the indicated proteins in whole-cell lysate or FLAG-immunoprecipitant from HEK293T cells transfected with *MARCHF6-*HA cDNA, along with an empty vector or *TXNDC15-*3xFLAG cDNAs. The cells were transfected with vectors expressing the indicated sgRNAs and pre-treated with arachidonic acid (50 µM for 4 hours) or BSA as a control. GAPDH was used as a loading control. (**H**) Immunoblots showing the indicated proteins in the membrane fraction (M) or the cytosolic fraction (C) from cells expressing the *ABHD2*-3xFLAG cDNA and the indicated sgRNAs. Cells were treated with MG132 (25 µM for 4 hours) or DMSO as a control.

Based on these analyses, we focused the functional dissection on the thioredoxin domain of TXNDC15. The thioredoxin domain harbors a single CXXS motif that is canonically associated with redox-active disulfide bonds and was shown to be involved in cilia biogenesis^12^. To test the potential role of redox catalysis in TXNDC15 function, we introduced the C220S mutation that abolished the redox-active thiol of TXNDC15. To our surprise, TXNDC15 (C220S) displayed an unabated function in promoting ABHD2 degradation (**Figure 3E**). To rule out the possible involvement of other reactive cysteine residues, we mutated all five cysteines to serine in TXNDC15 and remarkably, the TXNDC15(5xC-S) mutant is still able to restore ABHD2 degradation (**Figures 3F and S3B**). Given the unique oxidative environment of the ER lumen, we also quantified the redox state of cysteines on TXNDC15 using a maleimide-based probe that specifically labels the oxidized cysteine residues (**Figure S3C**). While this assay indicates the presence of oxidized cysteines on TXNDC15 at baseline (**Figure S3D**), mutating Cys220 to serine did not reduce the abundance of the oxidized species (**Figure S3E**). Together, these data suggest that TXNDC15 participates in ERAD via a non-canonical mechanism that relies on the non-catalytic activity of its thioredoxin domain.

### TXNDC15 promotes substrate exit from the endoplasmic reticulum

The surprisingly dispensable role of the catalytic activity of TXNDC15 prompted us to consider alternative mechanisms of action for its thioredoxin domain. Since thioredoxin domains may serve as interfaces that interact with numerous ER proteins^13^, we considered the possibility that TXNDC15 may function as an adaptor protein for recruiting MARCHF6 substrates. To test this possibility, we examined the interaction between ABHD2 and MARCHF6 in the presence or absence of TXNDC15. Surprisingly, the interaction between ABHD2 and MARCHF6 persisted even in the absence of TXNDC15 (**Figure 3G**), indicating that TXNDC15 is not required for recruiting substrate to MARCHF6. Rather, in the absence of TXNDC15, the substrate may stay trapped at the MARCHF6 complex due to failure of ubiquitination or translocation. Consistent with this prediction, TXNDC15 deficiency leads to the accumulation of ABHD2 in the membrane fraction that contains the predominant fraction of ER-localized proteins (**Figure 3H**).

The precise mechanism underlying functional coordination between TXNDC15 and MARCHF6 likely requires structural analysis. However, we reasoned that the thioredoxin domain may also interact with additional ER lumen proteins that assist MARCHF6 substrate degradation. Accordingly, we performed MS-based proteomics on immunoprecipitated, 3xFLAG-tagged full-length or thioredoxin domain-truncated TXNDC15 (**Figure S3F**). Crosslinkers were applied to enrich weak or transient interactions mediated specifically by the thioredoxin domain. Notably, we identified numerous proteins involved in the quality control of ER-lumen glycoprotein to be associated with TXNDC15 specifically via its thioredoxin domain (**Figure S3F and Data S3**) Among the most significantly enriched proteins that interact with full-length TXNDC15 is Glucosidase II Alpha Subunit GANAB, which trims the glucose residues from the immature glycans prior to either the maturation of glycoproteins or their retro-translocation and degradation. We reasoned that the glycan processing proteins may be recruited by TXNDC15 and act on ERAD substrates. To examine the functional connection between glycan processing to TXNDC15, we constructed a fusion protein in which the N-terminal portion, including the thioredoxin domain, of TXNDC15 is removed and spliced with GANAB (**Figure S3G**). We found that the fusion protein, when overexpressed, displayed a profound effect in promoting ABHD2 degradation and partially compensated for the absence of wild-type TXNDC15. Of note, although TXNDC15 itself is glycosylated (**Figure S3H**), introducing mutations that prevent the glycosylation does not abolish its function (**Figure S3I**). Collectively, these data support a model in which TXNDC15 assists the exit and degradation of MARCHF6 substrates from the ER, an activity likely cooperated by additional ER protein quality control factors, including the glycan processing machinery.

### TXNDC15 loss rewires the ER proteome and lipid metabolism

MARCHF6 has been shown to integrate numerous metabolic and environmental cues and functions as a hub for the coordination of multiple lipid metabolic pathways. We reasoned that, as an essential cofactor for ERAD via MARCHF6, TXNDC15 would play a significant role in shaping the ER proteome and the cellular lipid profiles. To profile localized changes in the ER proteome upon the perturbation of the MARCHF6-TXNDC15 complex, we employed a rapid immunopurification method that we developed to enrich ER membranes from cells expressing 3xHA-tagged EMC3 protein using anti-HA affinity beads (**Figure 4A**)^14^. Remarkably, we observed that numerous proteins that increased upon the ablation of *MARCHF6* also increased prominently in *TXNDC15*-knockout cells (**Figure 4B and Data S4**). Candidate MARCHF6 targets displayed significant enrichment among the proteins that accumulated in the ER under *TXNDC15* deficiency (**Figure 4C**), supporting their overlapping function in shaping the ER proteome.

**Figure 4.**
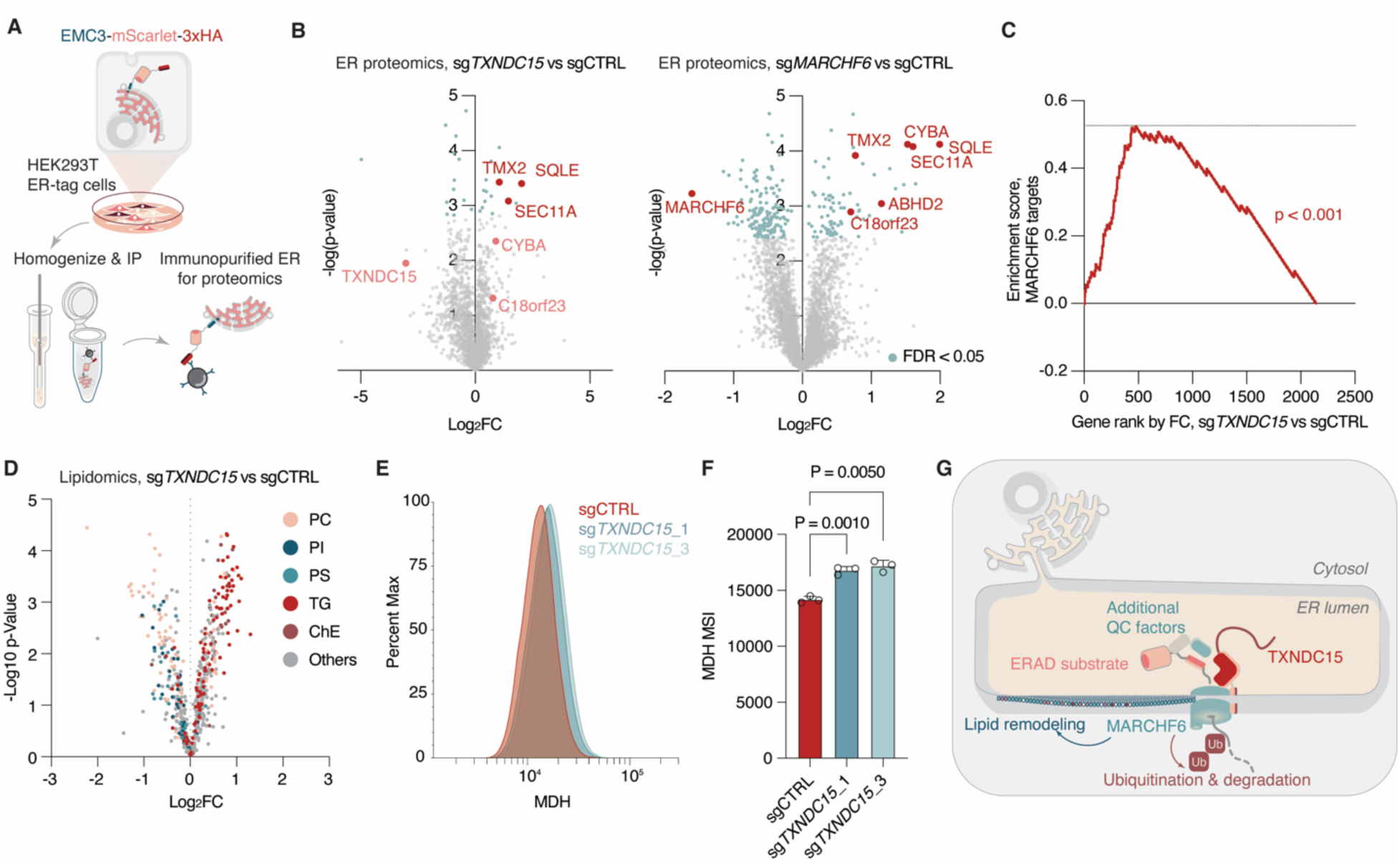
Dysfunction of TXNDC15 rewires the ER proteome and cellular lipid metabolism. (**A**) Schematic of the workflow for rapid immunopurification of endoplasmic reticulum. (**B**) Volcano plot showing the fold change and -log p-value of protein abundance in purified ER from cells expressing sgRNAs targeting *TXNDC15, MARCHF5* or intergenic control. (**C**) Enrichment analysis of candidate MARCHF6 targets (proteins showing significant accumulation upon *MARCHF6-*knockout with FDR < 0.05) among proteins displaying increased abundance in *TXNDC15-*knockout ER proteomics, as ranked by fold change. P-value computed by the permutation test of the data with 100,000 random shuffling of data labels. (**D**) Volcano plot showing the fold change and -log p-value of lipid abundance in purified ER from cells expressing sgRNAs targeting *TXNDC15* or intergenic control. Selected lipid classes are labeled in the indicated colors. (**E**) Histogram showing the signal intensity distribution of monodansylpentane-stained HEK293T cells expressing sgRNA against *TXNDC15* or a non-targeting control. (**F**) Mean fluorescence signal intensity (MSI) of monodansylpentane-stained HEK293T cells expressing sgRNA against *TXNDC15* or a non-targeting control. Data are mean ± s.d., representing three biologically independent samples. P-values were calculated from one-way ANOVA. (**G**) Schematic showing the proposed model of TXNDC15 function in mediating MARCHF6-dependent ERAD and lipid homeostasis

Given the pivotal role of MARCHF6 in modulating lipid synthesis activity in the ER^9^, we next performed lipidomic analysis on HEK293T cells expressing nontargeting or *TXNDC15* sgRNAs. Ablation of *TXNDC15* expression induced a significant rewiring of the global lipidomic profile. Specifically, we observed significant enrichment for multiple triglyceride (TG) and cholesterol-ester (ChE) subspecies and depletion of phosphatidylserine/phosphatidylinositol (PS/PI) species (**Figure 4D and Data S5**). This profile is reminiscent of lipidome remodeling in the absence of MARCHF6^15^. To validate the increased triglycerides seen in lipidomics, we stained sgCTRL and sg*TXNDC15* cells with a selective fluorophore for lipid droplet Monodansylpentane (MDH); *TXNDC15*_KO cells displayed a modest but significant increase in MDH fluorescence intensity in agreement with the accumulation of neutral lipid species (**Figures 4E and 4F**). Together, our results demonstrated that TXNDC15 is necessary for maintaining the cellular lipid homeostasis (**Figure 4G**).

## Discussion

The endoplasmic reticulum is a dynamic compartment for metabolism where multiple pathways for nutrient sensing and stress adaptation converge. The ER plays a particularly important role in the biosynthesis of numerous lipid species, which are critical for the physiological functions of membranes, cell signaling and energy homeostasis^16,17^. This process is also intricately intertwined with environmental parameters including oxygen levels, redox homeostasis and exogenous lipid availability. It is therefore not surprising that eukaryotic cells have evolved multiple nutrient and stress sensors residing on the ER that fine-tune lipid metabolism. The dynamic nature of ER proteome enables these nutrient-sensing pathways to co-opt protein post-translational processing, degradation and trafficking for a timely response to changes in cellular lipid profiles. Achieving targeted and coordinated remodeling of the ER proteome under these scenarios would require dedicated arrays of adaptors, effector and auxiliary proteins to recognize and process their targets. Our findings on the role of TXNDC15 in assisting MARHCF6 function highlight the complexity and knowledge gap in the molecular mechanisms of these processes.

MARCHF6 is a highly conserved protein that displays a swift response to membrane lipid saturation, a crucial physiological parameter that must be tightly regulated. Additional evidence suggests that MARCHF6 responds to a rather wide range of metabolic cues, including cholesterol and NADPH levels, consistent with its role as an integrator for cellular metabolic signals^18,19^. The molecular basis for MARCHF6 lipid sensing remains elusive, although recent structural studies on Doa10 highlighted some unusual features of the protein, including a large central pore filled with lipid and lipid-interacting residues essential for the ubiquitin-conjugating functions of Doa10^20,21^. Apart from its role as a lipid saturation sensor, Doa10/MARCHF6 displays a protein translocase activity in a reconstituted system^22^; how MARCHF6 coordinates its lipid sensing, ubiquitin conjugation and protein translocation functions requires further clarification, and our discovery of the role of lumen protein TXNDC15 suggests that additional layers of regulation are in play.

Modulation of lipid homeostasis via MARCHF6 exemplifies the complexity of post-translational regulation of metabolism in membrane-enclosed organelles^23^. Such modality of regulation enables localized sensing and rewiring of metabolism in these sub-compartments, and may therefore play an important role in maintaining metabolic homeostasis in heterogeneous organelle populations. Similar regulatory circuits have been found in both the ER and other organelles^8,24,25^, and the multitude of proteins with fast turnover suggests that more may be co-opted for regulatory purposes. Dissecting the molecular components of these additional pathways would provide insights into metabolic homeostasis at both the cellular and organismal levels.

## Acknowledgments

We thank all members of the Birsoy laboratory for helpful suggestions. Data was generated by the Proteomics Resource Center (RRID:SCR_017797) and Genomics Resource Center at The Rockefeller University.

## Funding

National Institutes of Health grant F99CA284249 (YL)

Human Frontiers Science Program Postdoctoral Fellowship (SL)

National Institutes of Health grant R01DK140337 (KB)

National Institutes of Health grant R01CA273233 (KB)

Mark Foundation Emerging Leader Award (KB)

Pew Innovation Fund Investigator (KB)

Chan Zuckerberg Biohub New York Investigator (KB)

## Author contributions

Conceptualization: YL and KB

Methodology: YL, SL, HA and KB

Investigation: YL, MYY, SL and HA

Funding acquisition: YL, SL and KB

Supervision: KB

Writing – original draft: YL

Writing – review & editing: YL and KB

## Data, code, and materials availability

All data are available in the main text or the supplementary materials.

## Supplementary Materials

Materials and Methods

Figs. S1 to S3

Data S1 to S5

## Supplementary materials

**Fig. S1.**
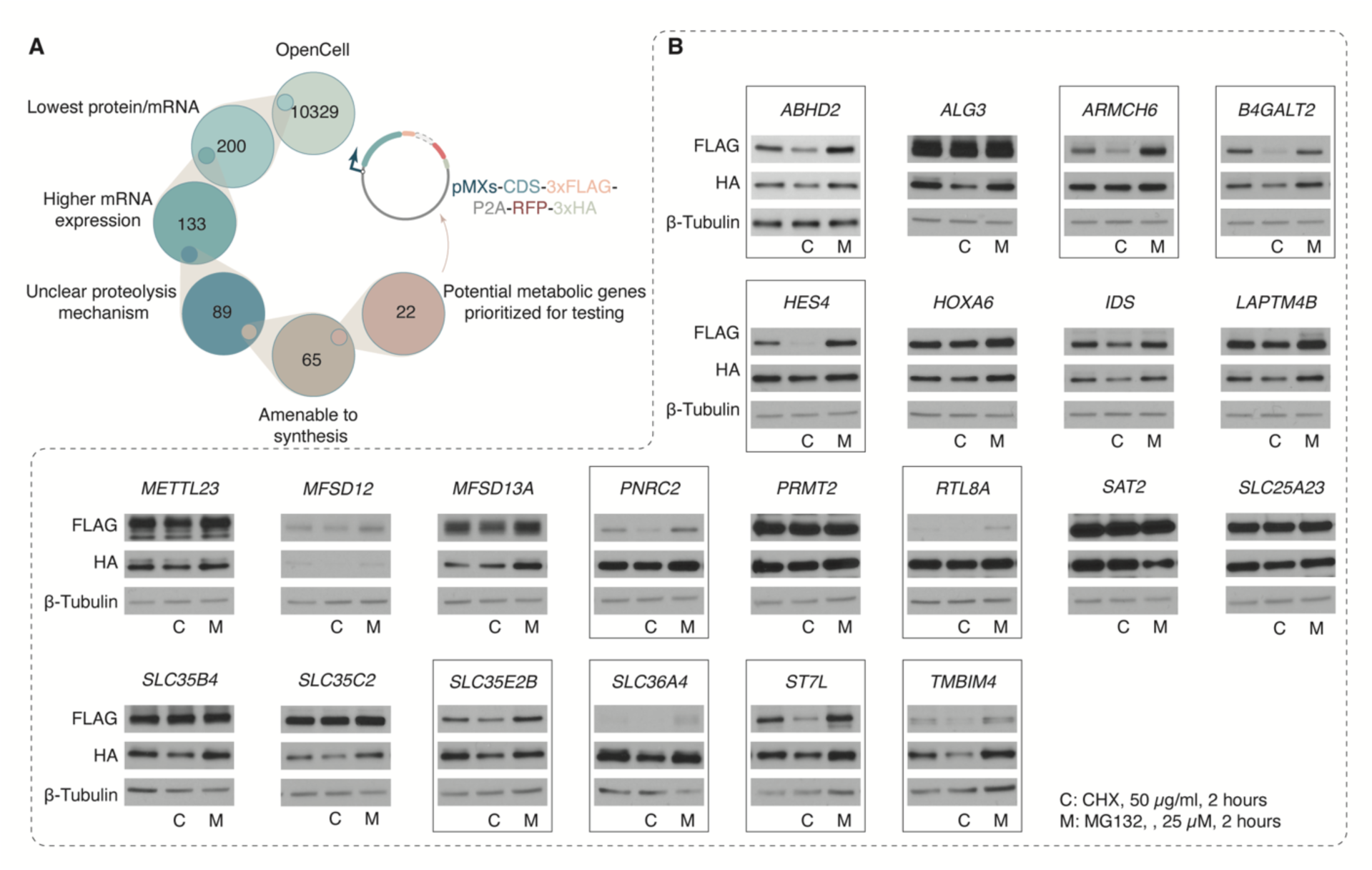
Schematic for the selection and validation of short-half-life protein candidates. (**A**) Diagram illustrating the selection and prioritization of short-half-life protein candidates from the *OpenCell* database, and the schematic of the reporter plasmid co-expressing the 3xFLAG-tagged coding sequence (CDS) of the candidate protein and 3xHA-tagged RFP from the same ORF, separated by a self-cleaving P2A peptide. (**B**) Immunoblots showing the indicated proteins from HEK293T cells transfected with reporter plasmids expressing the indicated short half-life candidate proteins. Cells were treated with cycloheximide (50 µg/ml), MG-132 (25 µM) or DMSO as a control for 2 hours prior to harvesting. β-tubulin was used as a loading control. Proteins showing behaviors consistent with proteasome-dependent degradation are highlighted by boxes.

**Fig. S2.**
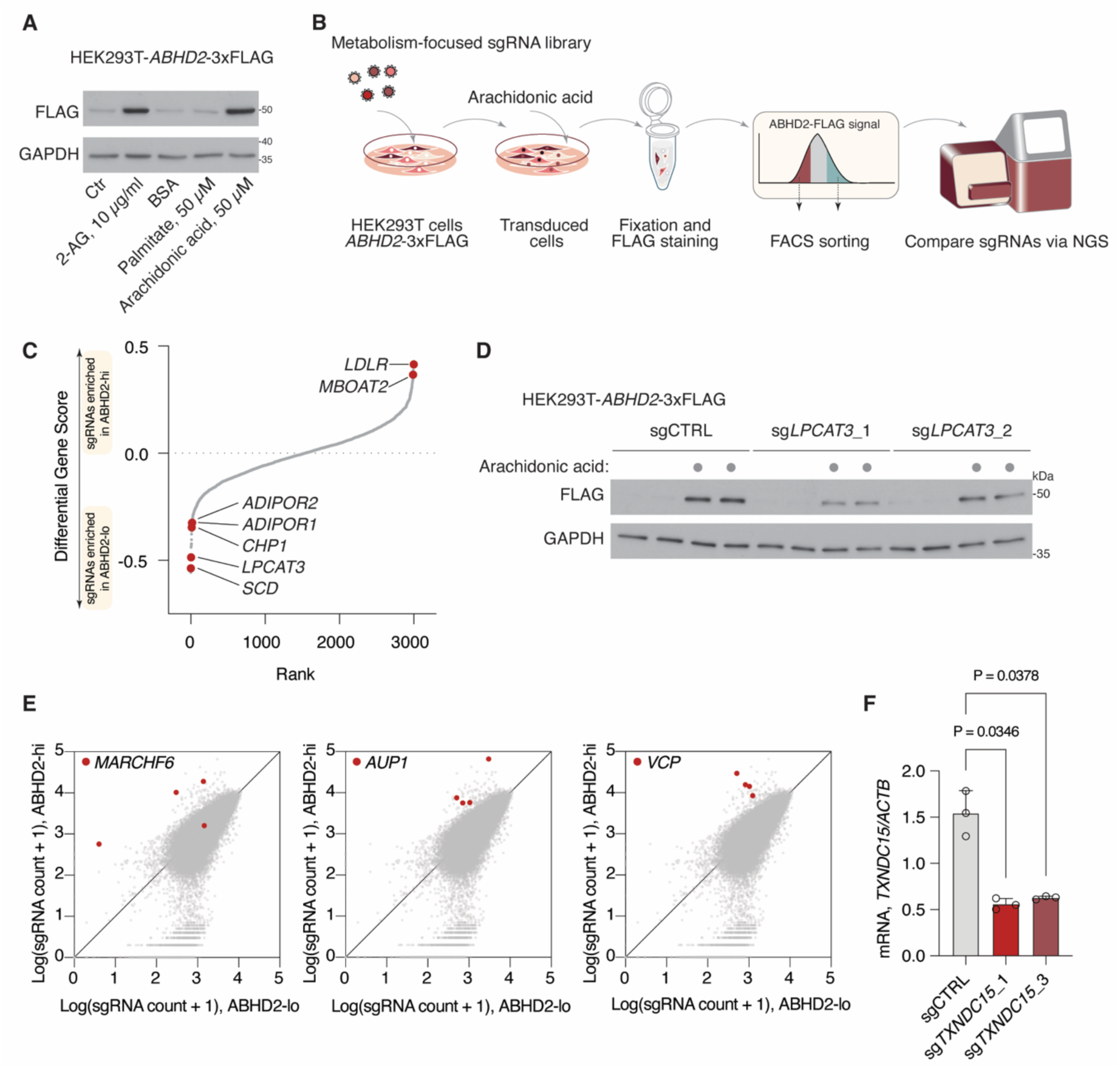
ABHD2 is degraded via the MARCHF6-dependent ERAD pathway in a lipid-sensitive manner. (**A**) Immunoblots showing the indicated proteins from HEK293T cells stably expressing ABHD2-3xFLAG, treated with the indicated lipid species for 4 hours or bovine serum albumin (BSA) as a control. β-tubulin was used as a loading control. (**B**) Schematic showing the workflow of the metabolism gene-focused, FACS-coupled CRISPR screen for regulators of ABHD2-3xFLAG stability. (**C**) Dot plot representing the CRISPR screen result. Points indicate the median fold change of sgRNA counts (gene score) of the quantified genes in the ABHD2-high versus the ABHD2-low fraction. (**D**) Immunoblots showing the indicated proteins from HEK293T cells stably expressing ABHD2-3xFLAG and transfected with plasmids expressing the sgRNAs targeting *LPCAT3* or intergenic control, treated with arachidonic acid or bovine serum albumin (BSA) as a control. GAPDH was used as a loading control. (**E**) Scatter plots showing the abundance of sgRNAs from the genome-wide CRISPR screen in the ABHD2-high versus the ABHD2-low fractions. sgRNAs targeting the indicated genes were highlighted in red. (**F**) Relative amount of *TXNDC15* mRNA in HEK293T cells transfected with plasmids expressing the indicated sgRNAs, as normalized by the mRNA levels of *ACTB*.

**Fig. S3.**
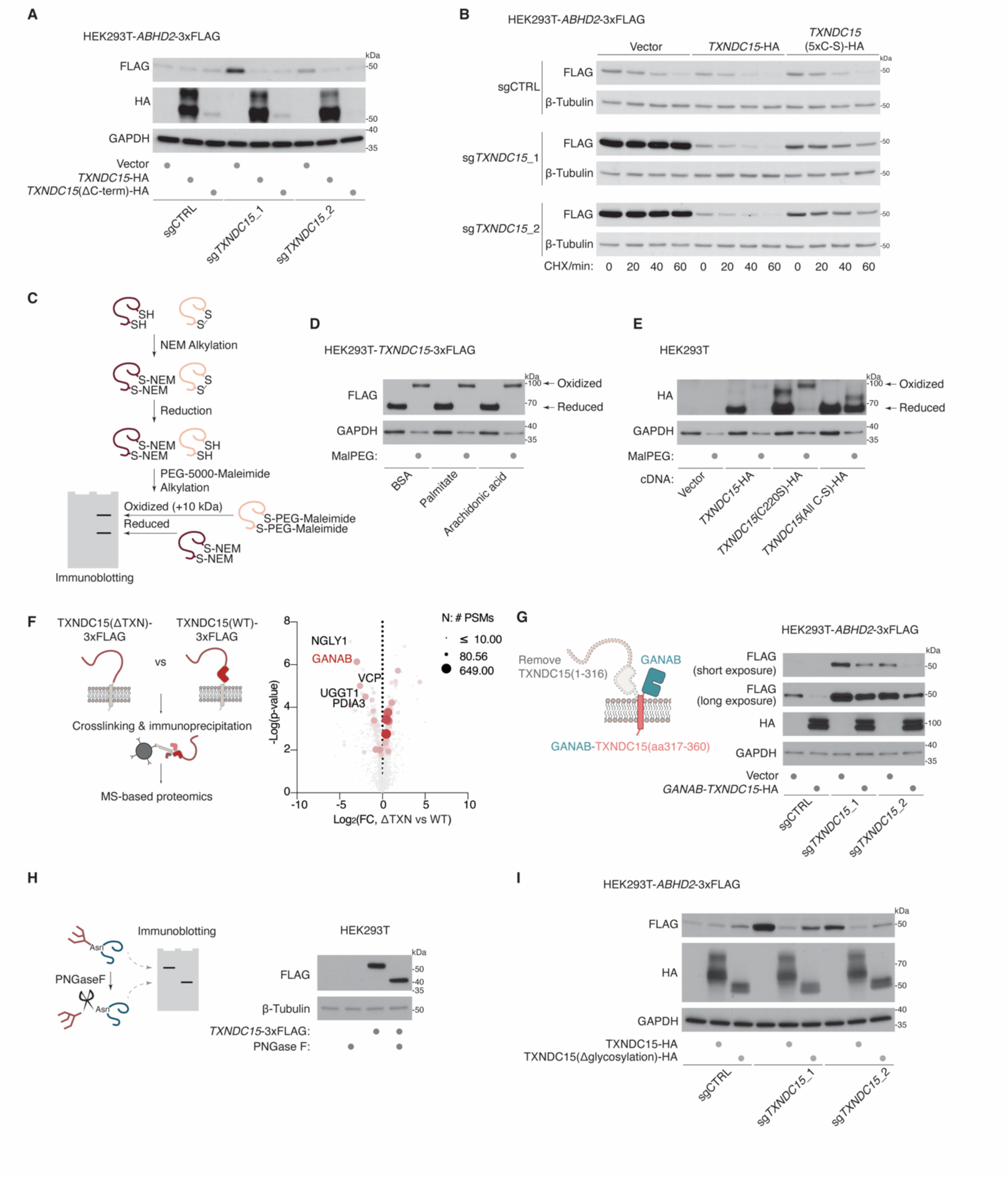
TXNDC15 promotes MARCHF6 substrates degradation via catalysis-independent mechanisms. (**A**) Immunoblots of the indicated proteins from HEK293T cells stably expressing *ABHD2*-3xFLAG cDNA and the indicated sgRNAs. Cells were complemented with cDNA of wild-type or the C-terminal truncated (1′aa350-360) mutant *TXNDC15*-HA. GAPDH was used as a loading control. (**B**) Degradation kinetics of ABHD2-3xFLAG as shown by immunoblots of the indicated proteins from HEK293T cells stably expressing *ABHD2*-3xFLAG cDNA, treated with cycloheximide (CHX, 50 µg/ml) or DMSO as a control for the indicated time. Cells were stably expressing the indicated sgRNAs and *TXNDC15-*HA, *TXNDC15*(5xC-S)-HA cDNA or an empty vector as a control. GAPDH was used as a loading control. (**C**) Schematic for the Maleimide-PEG5000 labeling assay for detecting cysteine oxidation. (**D**) Immunoblots of the indicated proteins from HEK293T cells stably expressing *TXNDC15*-3xFLAG cDNA, labeled with Maleimide-PEG5000 or N-ethylmaleimide as a control. Cells were treated with 50 µM of the indicated lipids for 8 hours. (**E**) Immunoblots of the indicated proteins from HEK293T cells stably expressing an empty vector or the indicated variants of *TXNDC15*-3xFLAG cDNA, labeled with Maleimide-PEG5000 or N-ethylmaleimide as a control. (**F**) Left, schematic of the TXNDC15 constructs used for immunoprecipitation-mass spectrometry-based proteomics analysis. Right, volcano plots showing the fold change and -Log10 p-value of protein abundance that co-immunoprecipitated with wild-type or thioredoxin-null TXNDC15 constructs. Bubble size and color indicate the summed peptide spectrum counts (Σ PSM) of the indicated protein. (**G**) Left, schematic of the wild-type TXNDC15 and GANAB-TXNDC15 chimeric constructs used in the experiment. Right, immunoblots of the indicated proteins from HEK293T cells stably expressing *ABHD2*-3xFLAG cDNA and the indicated sgRNAs. Cells were complemented with cDNA of wild-type *TXNDC15*-HA or chimeric *GANAB-TXNDC15*-HA. GAPDH was used as a loading control. (**H**) Left, schematic of the PNGaseF assay for detecting protein N-glycosylation. Right, Immunoblots of the indicated proteins from HEK293T expressing *TXNDC15*-3xFLAG cDNA or an empty vector as control. Protein lysate was treated with PNGaseF or mock-treated as a control. (**I**) Immunoblots of the indicated proteins from HEK293T cells stably expressing *ABHD2*-3xFLAG cDNA and the indicated sgRNAs. Cells were complemented with cDNA of wild-type or the glycosylation-defective (N194Q) mutant *TXNDC15*-HA. GAPDH was used as a loading control.

**Data S1. Selection of candidate short-half-life genes**

**Data S2. Results of all CRISPR screens**

**Data S3. IP-MS proteomics of TXNDC15 interaction partners, TXNDC15-WT vs TXNDC15(deltaTXN)Data S2. Results of all CRISPR screens**

**Data S4. ER-IP proteomics, sg*TXNDC15* or sg*MARCHF6* vs sgCTRL**

**Data S5 Lipidomics analysis for HEK293T cells, sgCTRL vs sg*TXNDC15***

