## Supplemental file for "Genetic dissection of rapid proteolysis identifies TXNDC15 as a key factor of ERAD and lipid homeostasis"

### Materials and Methods

#### Key Resource Table

| REAGENT or RESOURCE | SOURCE | IDENTIFIER |
|---|---|---|
| **Antibodies** | | |
| FLAG-M2 | Sigma | F1804 |
| beta-Tubulin | GeneTex | GTX101279 |
| Ubiquitin | Bethyl Laboratories | A300-317A-T |
| GAPDH | GeneTex | GTX627408 |
| HA | Cell Signaling Technology | 3724S |
| Calnexin | Cell Signaling Technology | 2679S |
| SQLE | Proteintech | 12544-1-AP |
| Anti-rabbit IgG, HRP-linked | Cell Signaling | 7074S |
| Anti-mouse IgG, HRP-linked | Cell Signaling | 7076S |
| Anti-Mouse Secondary Antibody, Alexa Fluor 568 | Thermo Fisher Scientific | A10037 |
| Anti-Rabbit Secondary Antibody, Alexa Fluor 488 | Thermo Fisher Scientific | A21206 |
| **Bacterial and Virus Strains** | | |
| NEB Stable Competent *E. coli* | NEB | C3040 |
| **Chemicals, Peptides, and Recombinant Proteins** | | |
| DMEM | GIBCO | 11965 |
| Trypsin | GIBCO | 25200 |
| BSA Fatty Acid-free | Alfa Aesar | J64944 |
| Palmitic acid | Cayman Chemical | 10006627 |
| Arachidonic acid | Cayman Chemical | 90010 |
| 2-arachidonyl glycerol | Cayman Chemical | 62160 |
| Penicillin-Streptomycin | GIBCO | 15140122 |
| FBS | Sigma | 12306C |
| N-ethylmaleimide | Sigma-Aldrich | E3876 |
| Cycloheximide | Cayman Chemical | 14126 |
| MG-132 | Selleck Chemicals | S2619 |
| Methoxy PEG Maleimide, 5000 | JenKem Technology | A3125-1 |
| Monodansylpentane | Abcepta | SM1000a |
| Phusion High-Fidelity PCR Master Mix with HF Buffer | NEB | M0531 |
| BsmBI | NEB | R0580 |
| T4 DNA Ligase | NEB | M0202 |
| X-tremeGENE 9 DNA Transfection Reagent | Roche | 6365779001 |
| BamHI | NEB | R3136 |
| NotI | NEB | R3189 |

| | | |
|---|---|---|
| PNGaseF | NEB | P0704S |
| Polybrene | Sigma | H9268 |
| Puromycin | Sigma | P8833 |
| Blasticidin | Invivogen | ant-bl |
| Dithiobis(succinimidyl propionate) | Thermo Fisher Scientific | 22585 |
| ANTI-FLAG(R) M2 MAGNETIC BEADS | Sigma-aldrich | M8823 |
| Pierce Anti-HA Magnetic Beads | Thermo Fisher Scientific | 88837 |
| **Critical Commercial Assays** | | |
| DNeasy Blood & Tissue Kit | QIAGEN | 69506 |
| Pierce BCA Protein Assay Kit | Thermo Fisher Scientific | 23225 |
| LunaScript RT SuperMix Kit | NEB | E3010L |
| RNeasy Mini Kit | QIAGEN | 74104 |
| Pierce Subcellular Protein Fractionation Kit | Thermo Scientific | 78840 |
| **Oligonucleotides** | | |
| Opti-sgCTRL | atcttgtggaaaggacgaaacaccgAACCTACGGGCTACGATACGgtttaagagctatgc | |
| Opti-sgTXNDC15.1 | atcttgtggaaaggacgaaacaccgAGAGTACAGTCACTACCGTTgtttaagagctatgc | |
| Opti-sgTXNDC15.2 | atcttgtggaaaggacgaaacaccgGAGCTCCTGCATGACCCGATgtttaagagctatgc | |
| Opti-sgTXNDC15.3 | atcttgtggaaaggacgaaacaccgCGGGGCAGAGAGTTAAAGTGgtttaagagctatgc | |
| Opti-sgMARCHF6.1 | atcttgtggaaaggacgaaacaccgTTGGCACTGCAATACGATATgtttaagagctatgc | |
| Opti-sgMARCHF6.2 | atcttgtggaaaggacgaaacaccgACATTGTATGGAAGAAAATTgtttaagagctatgc | |
| ACTB-qPCR-F | CATGTACGTTGCTATCCAGGC | |
| ACTB-qPCR-R | CTCCTTAATGTCACGCACGAT | |
| TXNDC15-qPCR-F | TGGTGCCGCTTTTCTGCCAGTT | |
| TXNDC15-qPCR-R | CAGCTACGGTGCCAAACCTGGT | |
| **Recombinant DNA** | | |
| CRISPR Cas9 metabolism sgRNA Library | (Birsoy et al., 2015) | N/A |
| CRISPR Cas9 E3ligase+ sgRNA Library | This study | N/A |
| CRISPR Cas9 Brunello genome-wide sgRNA Library | Addgene | 73179 |
| LentiCRISPRv2-Opti | Addgene | 163126 |
| pMXS-IRES-puro | Cell Biolabs | RTV-014 |
| pMXS-IRES-Blast | Cell Biolabs | RTV-016 |
| pLV-EF1a-IRES-Puro | Addgene | 85132 |
| pMXs-ABHD2-GGS-3xFLAG-P2A-RFP-IRES-BLAST | This study | N/A |

| | | |
|---|---|---|
| pMXs-ALG3-GGS-3xFLAG-P2A-RFP-IRES-BLAST | This study | N/A |
| pMXs-ARMCX6-GGS-3xFLAG-P2A-RFP-IRES-BLAST | This study | N/A |
| pMXs-B4GALT2-GGS-3xFLAG-P2A-RFP-IRES-BLAST | This study | N/A |
| pMXs-HES4-GGS-3xFLAG-P2A-RFP-IRES-BLAST | This study | N/A |
| pMXs-HOXA6-GGS-3xFLAG-P2A-RFP-IRES-BLAST | This study | N/A |
| pMXs-IDS-GGS-3xFLAG-P2A-RFP-IRES-BLAST | This study | N/A |
| pMXs-LAPTM4B-GGS-3xFLAG-P2A-RFP-IRES-BLAST | This study | N/A |
| pMXs-METTL23-GGS-3xFLAG-P2A-RFP-IRES-BLAST | This study | N/A |
| pMXs-MFSD12-GGS-3xFLAG-P2A-RFP-IRES-BLAST | This study | N/A |
| pMXs-MFSD13A-GGS-3xFLAG-P2A-RFP-IRES-BLAST | This study | N/A |
| pMXs-PNRC2-GGS-3xFLAG-P2A-RFP-IRES-BLAST | This study | N/A |
| pMXs-PRMT2-GGS-3xFLAG-P2A-RFP-IRES-BLAST | This study | N/A |
| pMXs-RTL8A-GGS-3xFLAG-P2A-RFP-IRES-BLAST | This study | N/A |
| pMXs-SAT2-GGS-3xFLAG-P2A-RFP-IRES-BLAST | This study | N/A |
| pMXs-SLC25A23-GGS-3xFLAG-P2A-RFP-IRES-BLAST | This study | N/A |
| pMXs-SLC35B4-GGS-3xFLAG-P2A-RFP-IRES-BLAST | This study | N/A |
| pMXs-SLC35C2-GGS-3xFLAG-P2A-RFP-IRES-BLAST | This study | N/A |
| pMXs-SLC35E2B-GGS-3xFLAG-P2A-RFP-IRES-BLAST | This study | N/A |
| pMXs-SLC36A4-GGS-3xFLAG-P2A-RFP-IRES-BLAST | This study | N/A |
| pMXs-ST7L-GGS-3xFLAG-P2A-RFP-IRES-BLAST | This study | N/A |
| pMXs-TMBIM4-GGS-3xFLAG-P2A-RFP-IRES-BLAST | This study | N/A |
| pLentiV2-OptiCas9deltaFLAG-sgCTRL-puro | This study | N/A |
| pLentiV2-OptiCas9deltaFLAG-sgMARCHF6.1-puro | This study | N/A |
| pLentiV2-OptiCas9deltaFLAG-sgMARCHF6.2-puro | This study | N/A |
| pLentiV2-OptiCas9deltaFLAG-sgTXNDC15.1-puro | This study | N/A |
| pLentiV2-OptiCas9deltaFLAG-sgTXNDC15.2-puro | This study | N/A |

| | | |
|---|---|---|
| pLentiV2-OptiCas9deltaFLAG-sgTXNDC15.3-puro | This study | N/A |
| pLV-TXNDC15-3xFLAG-blast | This study | N/A |
| pMXs-TXNDC15-3xFLAG-blast | This study | N/A |
| pMXs-TXNDC15(C220S)-3xFLAG-blast | This study | N/A |
| pMXs-TXNDC15(C220S)-3xFLAG-puro | This study | N/A |
| pMXs-TXNDC15-HA-puro | This study | N/A |
| pMXs-TXNDC15(C220S)-HA-puro | This study | N/A |
| pMXs-TXNDC15(NoCys)-HA-puro | This study | N/A |
| pMXs-TXNDC15(NoTxn)-HA-puro | This study | N/A |
| pMXs-TXNDC15(NoLoop)-HA-puro | This study | N/A |
| pMXs-TXNDC15-3xFLAG-blast | This study | N/A |
| pMXs-TXNDC15(delTXN)-3xFLAG-blast | This study | N/A |
| pMXs-TXNDC15(NoGly)-HA-puro | This study | N/A |
| pMXs-TXNDC15-OST4-3xFLAG-blast | This study | N/A |
| pMXs-TXNDC15-deltaCterm-HA-puro | This study | N/A |
| pCW57.1-TXNDC15-HA-puro | This study | N/A |
| pMXs-GANAB-TXNDC15-HA-puro | This study | N/A |
| pMXs-MARCHF6-HA-blast | This study | N/A |
| pMXs-EMC3-mScarlet-3xHA | Liu et al. 2026 | N/A |
| **Other** | | |
| Z2 Coulter Counter | Beckman | Model Z2 |
| SpectraMax Microplate Reader | Molecular Devices | Model M3 |
| Primovert Microscope | Carl Zeiss | 415510-1105-000 |
| REVOLVE4 Microscope | Echo Laboratories | FJSD1001 |
| Nikon A1R MP multiphoton confocal | Nikon | A1R |

#### Contact for Reagent and Resource Sharing

### Cell Lines, Compounds and Constructs

HEK293T cells were purchased from ATCC and authenticated by STR profiling. All cell strains were test routinely for mycoplasma contamination and were verified to be mycoplasma-free prior to experiments.

### Cell Culture Conditions

All cells were cultured in DMEM (GIBCO) with 4.5 g/L glucose, 110 mg/L pyruvate, 4 mM glutamine, and supplemented with 10% fetal bovine serum and 1% penicillin/streptomycin. All cells were kept in incubators set at 37°C, 21% O2 and 5% $CO_2$.

### Generation of overexpression and knockout constructs

Gene fragments encoding wild-type and mutant TXNDC15, MARCHF6, ABHD2 and other candidate unstable proteins were purchased from Twist Bioscience and Gibson-assembled into pMXs-IRES-blast or pMXs-IRES-puro vectors. Knockout vectors were constructed by assembling oligonucleotide (Integrated DNA Technologies) containing sgRNA spacer sequence and flanking nucleotides into LentiCRISPRv2-opti expression vector using the NEBuilder® HiFi DNA Assembly Master Mix. For initial screening of unstable proteins, a plasmid was constructed by cloning the BamHI-XhoI-3xFLAG-P2A-RFP-HA fragment into the pMXs-IRES-blast plasmid. The plasmid was then linearized with BamHI and XhoI, and gene fragments containing the sequences flanking these restriction sites and in frame with 3xFLAG-P2A-RFP-HA were Gibson-cloned into this expression vector.

### Generation of knockout and overexpression cell lines

Cells stably expressing *ABHD2*-3xFLAG. *MARCHF6*-HA, *TXNDC15*-3xFLAG, *TXNDC15*(ΔTXN)-3xFLAG and other *TXNDC15* mutants were constructed by retrovirus-mediated expression vectors. Briefly, pMXs expression plasmids were co-transfected with VSV-G and Gag-Pol plasmid to HEK293T cells using the XtremeGene9 transfection reagent. Media was changed 24 hours post-transfection and virus-containing media was harvested 48 hours post-transfection and cleared by passing through 0.45 µM syringe filters. The cells were transduced by incubation in the presence of virus and 4 µg/ml Polybrene and centrifugation at 2,200 rpm for 80 minutes on a Beckman Allegra X-14R Centrifuge with SX4750 rotor. For initial screening of candidate unstable protein and co-immunoprecipitation experiments, overexpression was achieved via transient transfection with XtremeGene9 transfection reagent. Knockout cell lines were generated by transfecting 1 µg LentiCRISPRv2-opti knockout vector to approximately 5E5 cells seeded in 6-well plates 24 hours prior to transfection. The cells were split into puromycin-containing media 24 hours post-transfection and selected for three days before puromycin removal and downstream analysis.

### FACS-based CRISPR screen

For the E3 ligase-focused CRISPR screen, a library was designed by combining human genes that encode proteins with annotated ubiquitin ligase activity from the Uniprot database and a predicted and curated list of E3 ligases and related proteins from the literature. The oligo pool was synthesized by Twist Bioscience, PCR-amplified, and Gibson-assembled into a modified LentiCRISPRv2-opti backbone (in which the FLAG sequence on Cas9 was removed) linearized with BsmBI. For the genome-wide CRISPR screen, sgRNA sequence and sgRNA scaffold were PCRed out from the liquid stock of human Brunello genome-wide sgRNA library and Gibson-assembled into a modified LentiCRISPRv2-opti backbone (in which the FLAG sequence on Cas9 was removed) linearized with BsmBI and EcoRI. The pooled plasmid was used to generate a lentivirus library in HEK293T cells. Virus titer was determined by infecting HEK293T cells with serially diluted virus and measuring cell viability after selection by puromycin.

HEK293T cells expressing ABHD2-3xFLAG was sorted into a 96-well plate as single cells and one clone with unimodal FLAG signal from flow cytometry was chosen for genetic screens. One day before infection, cells of approximately 1,000X the library size were seeded at a density of 5E5 per well in 6-well plates. Infection was done in the presence of 4ug/ml Polybrene and spin-infected by centrifugation at 2,200 rpm for 80 minutes on a Beckman Allegra X-14R Centrifuge with SX4750 rotor. The cells were expanded 1 day after spinfection, and puromycin selection was started 48 hours post-infection. After three days of selection, puromycin was removed and cells were allowed to grow for 2 days before harvested for FACS. For the whole-genome screen, cells were treated with 50 µM 18:1 lyso-phosphatidylethanolamine overnight before harvest. Cells were dissociated with FACS buffer (1% bovine serum albumin and 2mM EDTA in PBS) and washed twice with PBS and resuspended in PBS at a concentration of approximately 1E8 per ml. Ice-cold methanol was added drop-wise with gentle vortexing to a final concentration of 80%. The cells were incubated for 10 minutes at -20 °C and rehydrated by adding PBS to a final volume percentage of 50%. The cells were washed once more with PBS and blocked for one hour rotating at 4 °C with FACS buffer (1% bovine serum albumin and 2mM EDTA in PBS). They were then stained overnight with AlexaFluor 488-conjugated anti-FLAG-M2 antibody in FACS buffer, rotating at 4 °C. The cells were washed three times prior to sorting, and cells with the highest 2% and lowest 20% FLAG signal were sorted on a Sony MA900 sorter. Approximately 1,000X library size number of cells were preserved before sorting as input. Genomic DNA was extracted with Qiagen DNeasy Blood & Tissue Kit or QiAamp DNA Blood Midi Kit. sgRNA sequence was amplified by PCR, and the amplicon was gel-purified and sequenced on the Nextseq or MiSeq platform (Illumina). Gene score for E3 ligase-focused screen was calculated as media of log2-transformed fold change in the sorted fraction versus the pre-sorted input. The weighted gene score for the whole-genome screen was calculated as media of log2-transformed fold change in the sorted fraction versus the pre-sorted input, weighted by the number of reads in the input and sorted cells combined.

**Immunoblotting**

Unless otherwise stated, approximately 5E5 cells were seeded in 6-well plates one day before drug treatment. Alternatively, approximately 1 million trypsinized cells were pelleted, washed by resuspending in 1ml ice-cold PBS and spun down at 1,000g for 3 minutes. Cells were homogenized by sonicating in 200 µl Membrane Lysis Buffer (2% SDS, 0.1% CHAPS, 50mM Tris-Cl, pH 7.4, 150mM NaCl). Protein concentration was determined by Pierce BCA assay. Laemmli buffer was added to a final concentration of 20%, and samples were boiled at 95 °C for 10 minutes and resolved on Novex 10%- 20% Tris-Glycine Gels. The proteins were transferred to Millipore PVDF membranes, blocked for 30 minutes with 5% BSA in TBS with 0.1% Tween-20 (TBS-T), and incubated overnight at 4 °C with primary antibodies diluted in 5% BSA in TBS-T. The membranes were washed 3 times with TBS-T for 5 minutes each and incubated with HRP-conjugated secondary antibodies in TBS-T for 1 hour. After three TBS-T washes for 5 minutes each, the membranes were briefly rinsed with deionized water and developed with ECL Chemiluminescent detection system (Perkin Elmer LLC) and film exposure using Premium autoradiography Films (Thomas Scientific) on a SRX-101A Film Processor (Konica Minolta). For non-reducing immunoblotting, cells were washed with 1ml PBS with 100 mM N-ethylmaleimide prior to lysis. The lysis buffer was also supplemented with 100 mM N-ethylmaleimide and boiling was skipped prior to electrophoresis.

**Immunofluorescence**

Approximately 2E5 HEK293T cells expressing the gene of interest were seeded on a coverslip pre-coated with poly-D-lysine one day before the experiment in a 6-well plate. The cells on the coverslip were washed 3 times with PBS and fixed with 4% formaldehyde in PBS for 20 minutes at room temperature. The coverslips were then washed three times with PBS for 5 minutes each and the cells were permeabilized with 0.1% Triton X-100 in PBS for 30 minutes. The cells were washed further for three times with PBS for 5 minutes each and blocked with 10% BSA in PBS for 30 minutes. The coverslips were incubated overnight at 4°C with primary antibodies diluted in 10% BSA in PBS (1:500 for FLAG-M2 antibody and 1:200 for others). The coverslips were then washed three times with PBS for 5 minutes each and stained with secondary antibodies (1:500 diluted in 10% BSA in PBS) for 1 hour at room temperature in the dark. The coverslips were then washed three times with PBS and stained with 0.1 μg/ml DAPI in PBS for 10 minutes at room temperature. The coverslips were washed again with PBS for three times and mounted on a glass slide in Molecular Probes ProLong Gold Antifade Mountant, cured overnight and sealed with nail polish. Images were taken on Nikon A1R MP multiphoton confocal using the Nikon Plan Apo γ 60X/1.40 oil immersion objective.

**Immunoprecipitation**

HEK293T cells were seeded at a density of 5 million per 10-cm dish 2 days prior to the experiment. Plasmids containing cDNAs of the protein of interest were transfected 24 hours prior to harvest. Cells were harvested by dissociating in 10 ml ice-cold PBS and pelleted by centrifugation at 1,000g x 2 minutes. The cells were resuspended in 1ml PBS with 0.25mg (dithiobis(succinimidyl propionate)) and incubated on ice for 10 minutes. DSP was quenched with 100 μl 1M Tri-Cl pH 7.4, and the cells were pelleted by spinning at 1,000g x 2 minutes. One milliliter of lysis buffer (1% Triton X-100, 50mM Tris-Cl, pH 7.4, 150mM NaCl and 1 mM EDTA with cOmplete™ EDTA-free Protease Inhibitor) was added and the mixture was incubated for 20min on ice. The sample was centrifuged for 10 min at 13,000 g and 200 μl supernatant was taken as Input. The rest of the supernatant was added to 5 μl of Sigma anti-FLAG(R) M2 magnetic beads slurry (pre-washed 3 times with lysis buffer) and rotated for 3.5 hours at 4 °C. The beads were washed four times with 300 mM NaCl, 150 mM Tris-Cl, pH 7.4, 0.1% Tween-20 and eluted by boiling in 200 μl Membrane lysis buffer at 95 °C for 10 minutes. Laemmli buffer was added to a final concentration of 20% and samples were boiled at 95 °C for 10 minutes. Samples were analyzed by immunoblotting.

**Cell fractionation**

Cell fractionation was performed using the Pierce Subcellular Protein Fractionation Kit (Thermo Scientific 78840) according to the manufacturer's instruction. Briefly, approximately 1E7 HEK293T cells were harvested in ice-cold PBS and pelleted by centrifugation at 1,000 g for 2 min. Cells were washed once with cold PBS and lysed in 100 μl cytosol extraction buffer (CEB) with 1:100 protease inhibitor cocktail and incubated for 10 minutes on ice. Following centrifugation at 500 g for 5 minutes, 160 μl supernatant was taken as the cytosolic fraction. The remnant supernatant was removed and the membrane fraction was extracted with 200 μl Membrane Extraction Buffer (MEB) with 1:100 protease inhibitor cocktail. The cells were vortexed vigorously and incubated for 30 minutes on ice, followed by centrifugation at 5,000 g for 5 minutes; 160 μl supernatant was taken as the membrane fraction.

**PNGaseF assay**

PNGaseF (NEB) assay was performed according to the manufacturer's instructions. Briefly. Approximately 1E6 to 2E6 cells were resuspended in 200 μl PBS and homogenized by sonication. 45 μl lysate was mixed with 5 μl denaturing buffer and boiled at 100 °C for 10 minutes and chilled on ice. Quenching solution mix (10 μl NP-40, 10 μl PNGase Buffer 2, 30 μl water) was added, and the reaction was split into 2 parts. 2.5 μl PNGase F was added for the experimental sample and omitted in the negative control. The samples were incubated at 37°C for 1 hour. Laemmli buffer was added to a final concentration of 20% and samples were boiled at 95 °C for 10 minutes. Samples were analyzed by immunoblotting.

**Measurement of cysteine redox state**

Cysteine redox state was measured based on a mass shift assay similar to that described before [26]. Cells were washed with SHE buffer (250 mM Sucrose, 10 mM HEPES, 1 mM EDTA, pH 7.4) containing 100 mM NEM and lysed in SHE buffer containing 100 mM NEM and 1% SDS. The cell lysate was buffer-exchanged using Micro Bio-Spin 6 chromatography columns (Biorad 732-6221) to remove NEM. The flowthrough was reduced in the presence of 2.5 mM DTT and 1% SDS by incubating at 37°C for 10 minutes. Reduced cysteines were alkylated with 50 mM Methoxy PEG Maleimide, 5000 (JenKem

Technology, A3125-1). Proteins were acetone-precipitated and re-solubilized in TBS with 1 % SDS for immunoblotting.

**Crosslinking immunoprecipitation and quantitative proteomics**

Immunoprecipitation was performed as described above, except that after the final wash, the beads were washed one more time with PBS and frozen at -80 °C. Beads underwent on-bead digestion with Trypsin for 3 hours. Supernatant was extracted and then reduced/alkylated, and proteins were then digested with Lys-C and Trypsin overnight. Digestion reactions were stopped with neat TFA. Samples then underwent Solid Phase Extraction, prior to being analyzed by LCMS (70 min analytical gradient and separated using a 50 cm EASYSprayer column (C18 reversed phase)). The mass spectrometer (ASCEND) was operated in high resolution/high mass accuracy mode. Generated data was searched and quantified using ProteomeDiscoverer/Mascot. Data were queried against the human database concatenated with user submitted sequence, along with Trypsin and Lys-C sequences. To produce statistical data, we used a version of Proteome Discoverer (v3.0) that is able to do the LFQ statistical analysis. We used the human database as well as a contamination database to produce the data. The data was then taken into Perseus. The data was then log 2 transformed, filtered out to remove any proteins that are considered contaminants, and we then filtered to only include proteins that had produced values in 2 out of 3 replicates in at least one sample group. We then imputed any missing values in order to produce a PCA plot and to perform two-sample t-tests.

**Quantification of neutral lipid by flow cytometry**

Approximately 1 million trypsinized HEK293T cells were pelleted by centrifugation at 1,000g x 2 minutes, resuspended in serum-free DMEM containing 100 µM Monodansylpentane. The cells were pelleted again, washed twice with FACS buffer (1% bovine serum albumin and 2mM EDTA in PBS) and resuspended in 1 ml FACS buffer before analyzed on an Attune NxT Flow Cytometer.

**Lipidomics analysis**

Approximately 5E5 cells were seeded per well in 6-well plates in triplicate 24 hours prior to harvest. The cells were washed twice with 1ml ice-cold 0.9% NaCl and subjected to lysis in 250 µl 80% MeOH buffer, by vigorous vortexing. 800 µl tert-Butyl methyl ether was then added, followed by vigorous mixing via vortex. The samples were then added 200 µl to induce phase separation and subjected to vortexing at 4 °C for 10 minutes. The samples were centrifuged for 10 minutes at 14,000g. The upper layer was transferred to a new Eppendorf tube and dried under a constant flow of gaseous nitrogen for lipidomics analysis. The bottom layer liquid was removed, and the pellet was solubilized in 8M Urea + 2% SDS in PBS to quantify protein concentration by BCA assay for normalization.

*LC-MS/MS analysis for lipids*

Lipid extracts were reconstituted in 50 µl 2-propanol (ULC/MS-CC/SFC grade, >99.95%; Biosolve B.V), shaken for 15 min, centrifuged at 10.000 g for 10 min, and 40 µl of each individual sample transferred to a glass vial with a micro glass insert. Lipids were separated by reverse phase chromatography (Accucore C30 column; 150 mm x 2.1 mm 2.6 µM 150 Å, Thermo Fisher Scientific) using a Vanquish Horizon UHPLC system (Thermo Fisher Scientific, Germering, Germany) coupled on-line to an Orbitrap Exploris 240 mass spectrometer (Thermo Fisher Scientific, Bremen, Germany) equipped with a HESI source. Lipids were separated at a flow rate of 0.3 mL/min and a column temperature of 50°C by a gradient elution starting at 10% B increasing in 20 min to 80% B, followed by an increase to 95% B within 4 min, and further 3 min until 100% B is reached and kept for 5 min, followed by a decrease to 10%B within 0.1 min and equilibration at 10% B for 7.9 min. Eluent A consisted of acetonitrile:water (50:50, v/v, both ULC/MS-CC/SFC grade, Biosolve B.V.) and Eluent B of 2-propanol:acetonitrile:water (85:10:5, v/v/v), both containing 5 mM ammonium formate (MS grade, Sigma Aldrich) and 0.1% (v/v) formic acid (ULC/MS-CC/SFC grade, Biosolve B.V.).

Full MS scans were acquired in positive and negative ionization mode with the following settings: spray voltage 3500 V/– 2500 V, sheath gas – 40 arb units, aux gas – 10 arb units, sweep gas – 1 arb unit, ion transfer tube – 300°C, vaporizer temperature – 370°C, EASY-IC run-start, default charge state – 1, resolution at m/z 200 – 120.000, normalized AGC target – 100%, maximum injection time – 100 ms, RF lens – 35%. For positive mode a scan range from m/z 200-1000 was covered and in negative mode full scans were acquired from m/z 200 – 1000 (until min 20) and from m/z 600-1700 (min 20 to 40).

Data-dependent acquisition (DDA) scans were acquired based on a cycle time of 1.3 s, a resolution of 15.000 at m/z 200, isolation window – 1.2 m/z, normalized stepped collision energies –17, 27, 37%, AGC target – 100%, maximum injection time – 60 ms, dynamic exclusion for 6 s after two-times fragmentation within 6 s.

*Data analysis for lipids*

Acquired raw files were processed within in Lipostar2 (Version 2.1.5b1). Briefly, super sample filters retaining only lipids with isotopic patterns and MS/MS were applied and remaining MS/MS spectra searched against LIPID MAPS structural database (LMSD, downloaded 21.10.2023; addition of lipid standards using LipoStar DB manager). For identification, lipids with 3 and 4 stars were automatically approved and searched for matching adducts and in-source fragments. Adducts were clustered and only approved features remained. These proposed identifications were manually checked for fragment matches and peak integration. Kendrick mass defect plots were used to validate the obtained identifications after manual inspection to filter out further false positive identifications. The obtained data matrix was exported and quantification was performed in Microsoft Excel 2016 by dividing the peak area of the identified lipid by the peak area of the corresponding internal standard, multiplied by the concentration of the internal standard. Total pmol of each lipid were further normalized to the total protein amount obtained from BCA assay [pmol/µg]. Data were further filtered by keeping lipids quantified in at least in 75% of the analyzed replicates per condition. For each obtained lipidome, mol percent were calculated and used for the final evaluation.

**Rapid ER purification for proteomics analysis**

*Sample preparation*

ER was purified from HEK293T cells expressing EMC3-mScarlet-3xHA (ER-tag). In brief, cells cultured in 2 x 15 cm dish were collected and washed twice with cold saline (0.9% NaCl), scraped into 3 ml of cold KPBS, and pelleted via centrifugation at 1,500 g at 4°C for 2 min. Cells were resuspended in 1 ml KPBS, 10 µl of cells were transferred into 90 µl of 1% (v/v) Triton lysis buffer (50 mM Tris-Cl pH 7.4, 150mM NaCl, 1% Triton X-100, 1mM EDTA) supplemented with protease inhibitor cocktail for a whole-cell protein sample. With one set of 30 strokes and another set of 30 strokes, the remaining sample was homogenized using a 2-ml homogenizer. After centrifugation at 3,000 g for 10 min at 4 °C, the homogenate was incubated with 300 ml of KPBS pre-washed anti-HA magnetic beads (Thermo Scientific Pierce 88837) on a rotator shaker for 5 min at 4 °C. Beads were washed three times in cold KPBS, then lysed with 50 µl 1% triton buffer for protein extracts by vortex for 10 seconds followed by a rotator shaker for 10Lmin at 4 °C. After magnetically removing the beads, samples were spun down at 20,000g for 15 min to remove potential cellular debris or bead contamination.

*Digestion*

Samples were dried and dissolved in 8 M urea, 50 mM triethylammonium bicarbonate (TEAB) and 10 mM dithiothreitol (DTT), and disulfide bonds were reduced for 1 hour at room temperature. Alkylation was performed using iodoacetamide (IAA) for 1 hour at room temperature in the dark. Proteins were precipitated by Wessel/Flügge extraction and pellets were dissolved in 100 mM TEAB with endopeptidase LysC (2% w/w, enzyme/substrate) and incubated at 37 °C for 2-3Lhours. Sequencing-grade modified trypsin (2 % w/w, enzyme/substrate) was added, and digestion proceeded overnight.

*Labelling and fractionation*

Peptide solutions were labelled with 270 µg aliquots of TMTpro (Thermo Scientific) for 1 h at room temperature and subsequently quenched with hydroxylamine for 15 min. An aliquot from each sample was combined for a ratio check, according to which the samples were mixed. The pooled sample was purified using a high-capacity reverse-phase cartridge (Oasis HLB, Waters) and the eluate was fractionated using high pH reverse phase spin columns (Pierce) according to manufacturer specifications, yielding eight fractions.

*LC-MS/MS*

Fractionated peptides were analyzed using an Easy-nLC 1200 HPLC equipped with a 250 mm × 75 µm Easyspray column connected to a Fusion Lumos mass spectrometer (all Thermo Scientific) operating in synchronous precursor selection (SPS)-MS3 mode34 (10 SPS events). Solvent A was 0.1% formic acid in water and solvent B was 80% acetonitrile, 0.1% formic acid in water. Peptides from the ER immunoprecipitation were separated across a 90-min linear gradient and peptides from the whole-cell lysate were separated across a 120-min linear gradient going from 7 to 33% solvent B at 300 nl min−1. Precursors were fragmented by CID (35% CE) and MS2 ions were measured in the ion trap. MS2 ions were fragmented by HCD (65% CE) and MS3 reporter ions were measured in the orbitrap at 50K resolution. Data were analyzed using Skyline v.20.1.1.158.

*Data analysis*

Raw files were searched through Proteome Discoverer v.2.3 (Thermo Scientific) and spectra were queried against the human proteome using Sequest HT with a 1 % false discovery rate (FDR) applied. Oxidation of M was applied as a variable modification and carbamylation of C was applied as a static modification. A maximum isolation interference of 50% was allowed and 80% matching SPS ions were required. Protein abundance values were used for further statistical analysis. Subsequent statistical analysis was performed within the Perseus framework. All values were log2-transformed and normalized to the median intensity within each sample. An FDR-corrected t-test (adjusted p-value = 0.05) was used to test for significant differences between sample groups. Imputed protein intensity values and precalculated p-values were loaded into R for analysis and subcellular localizations were generated as described below. Volcano plots were produced by filtering to proteins with at least 20 peptide spectrum matches, with proteins defined as hits if they passed a log-fold change cutoff of 2.5 and an adjusted p-value cutoff of 0.05.

**Quantification and statistical analysis**

Statistic analysis, unless otherwise stated, were performed using built-in functions and statistic tools of Prism9 (Graphpad Software). The specific statistic tests and number of biological replicates were stated in the figure legends.
